# Precision-engineered biomimetics: the human fallopian tube

**DOI:** 10.1101/2023.06.06.543923

**Authors:** Ashleigh J. Crawford, André Forjaz, Isha Bhorkar, Triya Roy, David Schell, Vasco Queiroga, Kehan Ren, Donald Kramer, Joanna Bons, Wilson Huang, Gabriella C. Russo, Meng-Horng Lee, Birgit Schilling, Pei-Hsun Wu, Ie-Ming Shih, Tian-Li Wang, Ashley Kiemen, Denis Wirtz

## Abstract

The fallopian tube has an essential role in several physiological and pathological processes from pregnancy to ovarian cancer. However, there are no biologically relevant models to study its pathophysiology. The state-of-the-art organoid model has been compared to two-dimensional tissue sections and molecularly assessed providing only cursory analyses of the model’s accuracy. We developed a novel multi-compartment organoid model of the human fallopian tube that was meticulously tuned to reflect the compartmentalization and heterogeneity of the tissue’s composition. We validated this organoid’s molecular expression patterns, cilia-driven transport function, and structural accuracy through a highly iterative platform wherein organoids are compared to a three-dimensional, single-cell resolution reference map of a healthy, transplantation-quality human fallopian tube. This organoid model was precision-engineered to match the human microanatomy.

**One sentence summary:** Tunable organoid modeling and CODA architectural quantification in tandem help design a tissue-validated organoid model.

## Introduction

The fallopian tubes (oviduct, uterine tube, salpinges) are a pair of muscular tube-like organs that are often overlooked in basic research. Their primary functions are to catch ovulating oocytes from the adjacent ovaries, provide an environment that supports survival of sperm and eggs for fertilization, and transport the developing embryo from the ovary to the uterus for implantation^1,2^. This process is achieved through peristalsis driven by the fallopian tube smooth muscle and motile cilia along the fallopian tube epithelium^1,2^. Structural or functional abnormalities are responsible for tubal pregnancy, a life-threatening condition if untreated^3,4^. Fallopian tubes also connect the uterine cavity and the peritoneal cavity, and serve as the anatomic structure for retrograde menstruation, one of the mechanisms of endometriosis which affects 10% of reproductive age women^5,6^. Recently, the fallopian tubes have received significant research attention because of the new paradigm in ovarian high-grade serous carcinoma (HGSC), the most common type of ovarian cancer. This paradigm posits that HGSCs arise from their precursor lesions, called serous tubal intraepithelial carcinomas (STICs), in the fallopian tube epithelium^7,8^. The exact mechanical and molecular mechanisms that govern these conditions in the fallopian tubes remain largely unknown, in part due to the lack of appropriate preclinical models^9,10^. HGSC and endometriosis, for example, do not spontaneously occur in mice as they do in humans^11–17^, so the field has turned to engineered mouse models and *in vitro* organoid models to study these human diseases in an environment with the highest physiological similarity^10,18,19^.

Histologically, the fallopian tube mucosa, muscularis and serosa regions are composed of unique combinations of epithelial cells, stromal and/or smooth muscle cells^20^; the extracellular matrix (ECM) of these regions varies as well. Beyond this compositional complexity, the organ also undergoes several dynamic changes in the body due to peritoneal fluid^21^, hormones, and follicular fluid that are released at various stages of the female reproductive cycle^22–24^. Each of these features and interactions are characteristic of a healthy fallopian tube.

To our knowledge, direct, three-dimensional, and quantitative comparison of organoid to organ microarchitecture and cellular composition has never been performed, although the field of organoid modeling has recognized the importance of validation through other tissue-omics-based methods^25,26^. Established fallopian tube organoid models typically culture a single cell type (epithelial cells) in a single ECM composition^27–29^, thus inherently limiting their anatomical similarity to the heterogeneous organ of origin. Validation is commonly performed through a rather cursory comparison of organoid architecture and molecular expression patterns to two-dimensional (2D) histological sections of reference human samples. Such comparisons only provide a cursory comparison and neglect the quantitative heterogeneity of tissues that is only revealed in 3D^30^.

Here, we present a novel, multi-compartment organoid model^31^ of the human fallopian tube that is highly tunable in both ECM and cellular composition of each compartment. This model facilitates spatial adjustment of organoid parameters to improve the correlation between organoid and tissue architecture. We compared this organoid model to expression patterns observed in histological tissue sections and observe a physiological response to menstrual cycle hormone stimulation, according to previously published validation parameters for fallopian tube epithelial organoids ^27–29^. In a novel functional assay, we evaluated the multi-compartment organoid’s functional capacity for cilia-driven transport of oocyte-mimicking beads along the lumen-facing epithelium. Unlike previous organoid models of the fallopian tube or other biological systems, we rigorously compared our organoid to a whole, transplantation-quality human fallopian tube using CODA^30^. CODA is a novel 3D pathology approach capable of quantitatively mapping the microanatomy of organs at cellular resolution. CODA has been used for 3D spatial mapping of normal and cancer-containing pancreas, liver, lung, breast, prostate, and skin tissues ^30,32–36^. In the human fallopian tube, CODA can label epithelial cells, stroma, nerves, vasculature, and additional structures at cellular resolution. We identified four biomimetic parameters, which were fine-tuned via side-by-side comparison of several iterations of the organoid model to a healthy fallopian tube reference map. This process enables for the first time a quantitative metric to measure the architectural and functional accuracy of an organoid model.

Through this quantitative iterative process, we produced a precision-engineered and functional organoid model of a human fallopian tube. This organoid model constitutes the basis for future studies investigating disease states of the fallopian tubes.

## Results

### Multi-compartment organoids self-organize to mimic the mucosal folds of the human fallopian tube

Our multi-compartment fallopian tube organoid was designed to mimic the *in situ* microenvironment where epithelial cells that line the fallopian tube lumen are supported by a thin basement membrane and surrounded by a collagen-rich stromal matrix (Fig. 1A). We captured this multi-extracellular matrix (ECM) microenvironment *in vitro* by first suspending fallopian tube epithelial cells (FTECs, 1 × 10^4^ cells/µL) in a small (1 µL) droplet of Matrigel, a recombinant basement membrane (Fig. 1B,C). This inner organoid core was embedded in a larger (10 µL total organoid volume) corona of collagen I (Fig. 1B,C). This protocol was based on an oil-in-water droplet microtechnology recently developed to generate multi-compartment tumor organoids^31^. Here, we leveraged the versatility of this approach by adjusting the concentrations of cells and ECM in each core and corona compartment to best fit the fallopian tube tissue architecture. We cultured FTECs (core) in a growth factor reduced Matrigel core and a 2-6 mg/mL collagen I corona. Compared to the standard fallopian tube organoid technique^27,28^, where FTECs (∼400 cells/µL) were suspended in 50 µL drops of Matrigel and supplemented with inhibitors/growth factors to encourage 3D growth (Fig. 1B,D), the multi-compartment model greatly improved the complexity of the extracellular environment and the resulting complex architecture of the epithelium. This new approach reduced the reagents required to establish organoid culture from 50 µL (standard) to 10 µL (multi-compartment), while producing much larger epithelial structures than the organoids produced by standard protocols. Although fewer reagents were required for each organoid, the cell-free regions in the multi-compartment organoids were closer in size to the fallopian tube lumen they resemble than the lumen-like region of standard organoids (Fig. 1E).

**Fig. 1.**
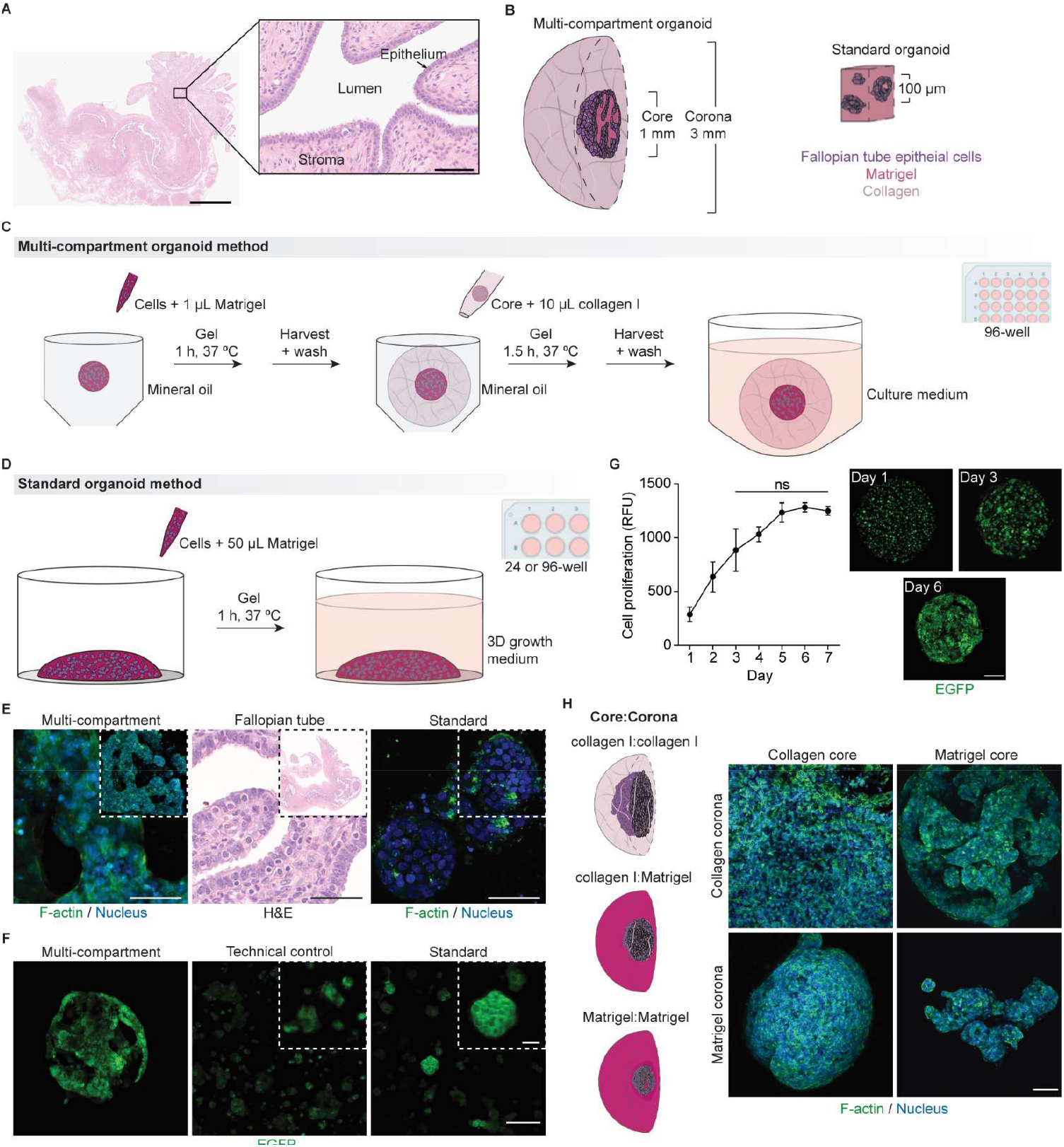
Human fallopian tube organoids. **A**, A healthy human fallopian tube tissue section. Hematoxylin and eosin (H&E): nuclei (purple) and ECM (pink). Scale bar, 5 mm. Zoom, 75 µm. Cartoons depicting **B**, mature fallopian tube organoids and the protocols to generate **C**, multi-compartment and **D**, standard fallopian tube organoids. **E**, Side-by-side comparison of multi-compartment organoids (left), human fallopian tube tissue (middle) and standard organoids (right). Inset shows one whole organoid or fallopian tube. F-actin (green) and nuclear DNA (blue). Fallopian tube tissue is an H&E stained tissue section. Scale bars, 50 µm. **F**, Organoids made with EGFP-tagged FTECs (green) on day 6 grown in the multi-compartment model (left), a technical control (middle), and the standard organoid model (right). Scale bar, 250 µm. Inset, 50 µm. **G**, PrestoBlue proliferation in multi-compartment organoids. Inset images of EGFP-tagged multi-compartment organoids (green). Scale bar, 250 µm. N = 3, n = 4+. Statistical test: one-way ANOVA, ns P > 0.05. Data are mean ± SEM. **H**, Representative images of multi-compartment organoids with all permutations to the organoid ECM assembly. F-actin (green) and nuclear DNA (blue). Scale bar, 100 µm. All organoid images are maximum intensity projections of stacks of confocal microscopy images.

Organoids grown in the multi-compartment model spontaneously assembled into intricate structures that mimic the tortuosity of the human fallopian tube’s mucosal folds (Fig. 1F, Fig. S1A). Spontaneous mucosal-like organization of primary FTEs, immortalized FTEs, and a mixture of FTE cell types was supported in the multi-compartment model (Fig. S1B,C). This architecture only formed when FTECs were cultured in the spherical cores of the multi-compartment model, and did not form when the same cells (1 × 10^4^ cells/µL) and culture conditions (unmodified cell culture medium) were plated according to the standard organoid generation technique (technical control Fig. 1F, Fig. S1D). In the standard model, organoids grew from single cells^27^, although not all FTEs developed into standard organoids (Fig. S1E). Multiple standard organoids often stacked on top of or next to each other, allowing (non-physiological) long-range interactions between separate organoids within the same culture well (Fig. 1F, Fig. S1E,F). These standard organoids assembled into small spherical clusters of cells with a necrotic core that did not reflect the complexity of the tissue architecture.

We note that the spontaneous assembly of FTECs into the mucosal architecture was not driven by cell proliferation. The architecture did not begin to assemble until after day 3, from which point there was no statistically significant change in proliferative activity (Fig. 1G). The formation of these folds therefore depends on cell re-organization and interactions with the immediate and surrounding ECM (Fig. 1H, Fig. S1G). While the ECM in each compartment can be adjusted to reflect any matrix-related changes in studies of fallopian tube-related diseases^37,38^, the intricate architecture resembling the fallopian tube’s lumen only formed when the matrix components were organized to resemble the fallopian tube anatomy, i.e., epithelial cells were embedded in a basement membrane (Matrigel) and surrounded by a collagen I rich environment. Unlike an (non-physiological) environment where cells were embedded in and surrounded by only collagen I, the Matrigel:collagen I core:corona combination (Fig. 1H) restrained the FTECs to a central mucosal region and provided the extracellular rigidity and structure necessary for organized folds to form. In the collagen I:Matrigel organoid, growth was contained within the core, but disorganized, and the Matrigel:Matrigel combination did not allow sufficient cell growth for the organoid architecture to mature.

These multi-compartment organoids can be further tuned to match the *in situ* microenvironment, such as by adding a third compartment containing smooth muscle cells. Here, we found that the dual-compartment model was sufficient for the spontaneous organization of an organoid architecture resembling the fallopian tube’s mucosal folds.

### Multi-compartment organoids maintain in situ protein expression and cellular structure

In addition to the similarity between the fallopian tube’s lumen and the multi-compartment organoid’s organization, the molecular expression patterns observed in these organoids also matched the tissue expression patterns according to previous FTE organoid validation parameters^27^. In the organoid, E-cadherin was localized to the lateral membranes of cells where strong adherens junctions form among the epithelium (Fig. 2A, Fig S2A). Ki-67-positive cells were scattered throughout the epithelium. Most FTECs were PAX8 positive and maintained a secretory status^27,39–42^. In this healthy fallopian tube organoid model, MUC16 staining was faint in some clustered regions. We expect an increase in MUC16 expression in future studies using this model investigating disease states of the fallopian tube^43,44^, for example, but limited MUC16 staining in this healthy baseline model aligns with our anticipated results.

**Fig. 2.**
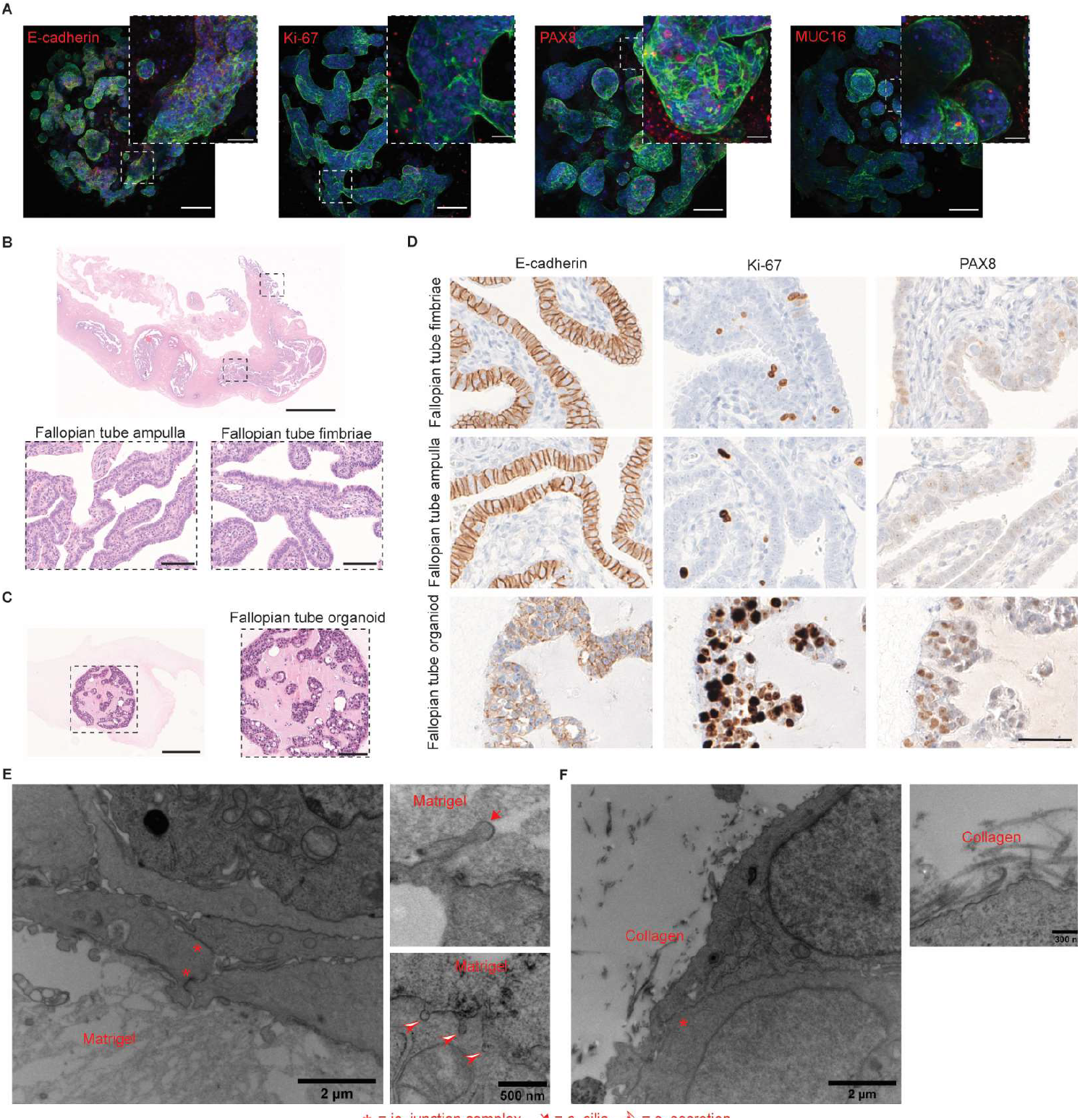
Fallopian tube organoids maintain tissue expression patterns and multi-cellular structure. **A**, Localized protein expression imaged via immunofluorescence in multi-compartment fallopian tube organoids. Images of organoids are maximum intensity projections of stacks of confocal microscopy images. Scale bar, 100 µm. Zoom, 20 µm. F-actin (green) and nuclear DNA (blue). Protein of interest in red, from left to right: E-cadherin, Ki-67, PAX8, MUC16. **B**, H&E image of a fallopian tube tissue section. Nuclei (purple) and ECM (pink). Scale bar, 5 mm. Left zoom is the fallopian tube ampulla, and right zoom is the fallopian tube fimbriae. Scale bar, 100 µm. **C**, H&E image of a fallopian tube organoid (multi-compartment). Scale bar, 300 µm. Zoom is the epithelial region of the organoid. Scale bar, 100 µm. **D**, IHC of tissue sections of the fallopian tube fimbriae (top), fallopian tube ampulla (middle) and fallopian tube organoid (bottom). From left to right: E-cadherin, Ki-67, PAX8. Scale bar, 50 µm. TEM of the **E**, cell-Matrigel interface and **F**, cell-collagen I interface. Scale bars, 2 µm. Zoom, **E**, 500 nm and **F**, 300 nm. Asterisk = junction complex, arrow = cilia, arrowhead = secretion.

There are slight architectural and cellular differences between the different regions of the fallopian tube. The percentage of epithelial cells at the fimbriated end that are ciliated, for example, is higher than throughout the ampulla^45–47^. However, the general mucosal folding architecture, ECM arrangement and epithelial/stromal cell types are preserved. We compared the staining patterns of select fallopian tube markers in the transplantation-quality, healthy fallopian tube fimbriae and ampulla (Fig. 2B) to those observed in our multi-compartment organoids (Fig. 2C) via immunohistochemistry (IHC) of FFPE tissue sections and found high similarity between the organoid and the tissue (Fig. 2D, Fig. S2B). We observed matching localization of E-cadherin to the lateral cell membrane in both the organ and the organoid. Ki-67-positive cells were dispersed among the epithelium of the organ and organoid, as well. In both cases, secretory cells (PAX8-positive) lined the epithelium; however, a larger number of PAX8-negative, ciliated cells existed in the tissue epithelium. Without any hormone stimulation, most cells in the organoid maintained a secretory status. We also confirmed that the multi-compartment organoid model formed adherens junctions (E-cadherin)^48,49^ and tight junctions (occludin)^50^, as was also observed in the standard organoid model via global protein expression (Fig. S2C). Compared to FTECs grown in standard 2D culture, E-cadherin expression was significantly increased in both 3D models.

Although occludin expression decreased slightly in organoid models compared to 2D cell cultures (Fig. S2C), the formation of tight junctions in multi-compartment and standard organoids was confirmed by transmission electron microscopy (TEM). Junction complexes (Fig. 2E, asterisks) formed between ciliated (Fig. 2E, arrow top right panel) and secretory (Fig. 2E, arrowheads bottom right panel) cells along the multi-compartment cell-Matrigel interface which simulates the cell-lumen interface of the tissue. At the cell-collagen interface, which simulates the tissue’s FTEC-stroma interface, junction complexes were still observed (Fig. 2F, asterisk); however, cells were not actively ciliated or secreting (right panel). These patterns in cell structure again reflected the tissue^27^ and improved the organ-level similarity observed in standard organoids. Standard organoids also formed tight junctions (Fig. S2D, asterisk), and showed both cilia (Fig. S2D, arrows) and secretions (Fig. S2D, arrowhead); however, in standard organoids, cilia formed on both the organoid interior (lumen-like region) and exterior (stroma-like region). In multi-compartment organoids, the cells adopted a more organized structural pattern based on the cell-lumen and cell-stroma simulated interfaces, and cells were polarized such that cilia faced the organoid lumen-like region.

### Activated multi-compartment organoids reproduce the organ’s function

The fallopian tubes have key roles in the female reproductive system, and as a result the organs are exposed to female hormones during the menstrual cycle. The steroid hormones estrogen and progesterone elicit morphological and functional responses^51^. Thus, we stimulated fallopian tube organoids with β-estradiol and progesterone, and their molecular response was measured via RT-qPCR (Fig. 3A, Fig. S3A). We observed overall increases in estrogen receptor 1 (ESR1) expression with β-estradiol treatment and a slight increase with progesterone treatment, matching previously reported responses in fallopian tube tissue^52^ and mouse FTEC standard organoids^28^. In the follicular phase when estrogen is high and progesterone is low^53^, progesterone receptor levels increase in the fallopian tube epithelium^54^ and, accordingly, progesterone receptor (PGR) expression increased when our organoids were treated with β-estradiol.

**Fig. 3.**
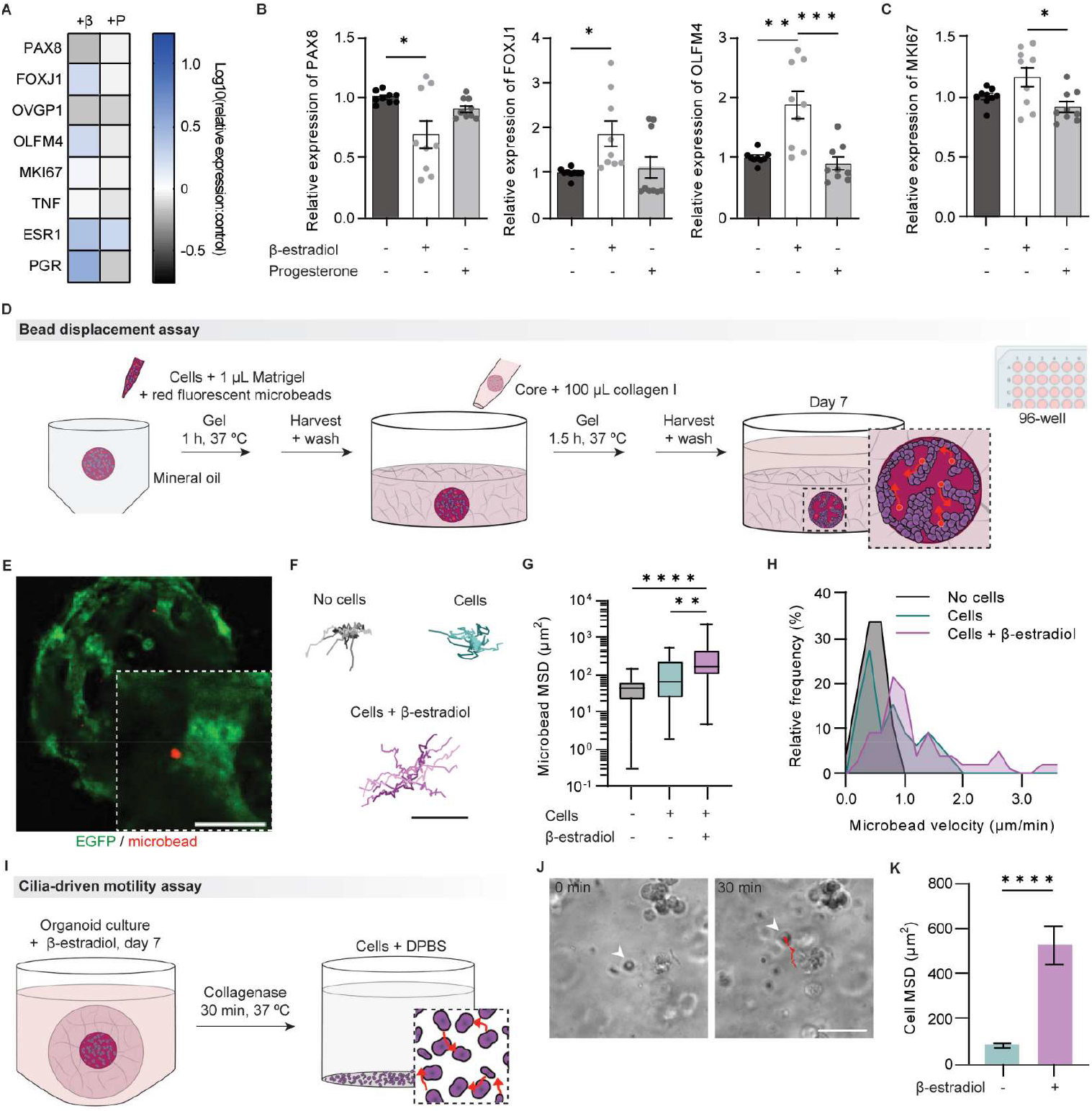
Multi-compartment organoids demonstrate functional similarity to the fallopian tube. **A**, RT-qPCR of β-estradiol (β) or progesterone (P) treated multi-compartment organoids. Data are log10(relative expression:control). Relative expression of genes related to **B**, ciliated differentiation and **C**, proliferation. **A-C**, N = 3, n = 3. **D**, Cartoon depicting bead displacement assay organoid generation. Arrows indicate microbead movement. **E**, A multi-compartment organoid used in the bead displacement assay with microbeads at the organoid’s epithelial-lumen interface. EGFP-tagged FTECs (green) and microbeads (red). Scale bar, 100 µm. **F**, Representative microbead trajectories from organoid cores with microbeads only (noise, black), microbeads with FTECs (teal), and microbeads with FTECs treated with β-estradiol (purple). Scale bar, 10 µm. **G**, Microbead MSDs over 8 h. **H**, Relative frequency of microbead velocities. **F**-**H**, N = 3, n = 4-46. **I**, Cartoon depicting cilia motility assay with organoid digestion. Arrows indicate cell movement in DPBS. **J**, Phase-contrast images from the cilia-driven motility assay. Cell trajectory (red). Scale bar, 50 µm. **K**, Cell MSDs in DPBS after organoid digestion. N = 3, n > 100. Statistical test: **B**-**C** and **G** one-way ANOVA, **K** unpaired t-test, ****P ≤ 0.0001, ***P ≤ 0.001, **P ≤ 0.01, *P ≤ 0.05. All data are mean ± SEM.

In addition to receptor level responses, β-estradiol treatment resulted in ciliated FTEC differentiation in multi-compartment organoids as the relative expression of secretory cell marker PAX8 decreased (Fig. 3B). Ciliated differentiation of epithelial cells in the fallopian tube and other gynecologic organs has been previously reported as a result of estrogen stimulation^26,55–57^. The relative expression of markers related to cilia motility/regulatory transcription factor (FOXJ1)^58,59^ and stem cell differentiation into ciliated cells (OLFM4)^60,61^ increased with β-estradiol stimulation. MKI67 relative expression was reduced when organoids were treated with progesterone compared to β-estradiol treatment (Fig. 3C). This decrease in proliferation when treated with progesterone aligns with progesterone’s ability to clear cells that have developed p53 mutations, such as HGSC precursor lesions^62–66^. This result demonstrated an appropriate response to progesterone stimulation of the organoid. Standard organoids did not always produce these expected responses to hormone treatment. MKI67 relative expression was reduced when treated with progesterone. However, markers of morphological cell type (PAX8, FOXJ1) were unchanged with β-estradiol stimulation, and instead there was a shift toward ciliated cells with progesterone treatment (Fig. S3B). Progesterone also prompted an unexpected increase in TNF expression.

From a functional perspective, the differentiation of FTECs into ciliated epithelial cells in response to β-estradiol stimulation is significant. While cilia and smooth muscle both are involved in the transport of an oocyte through the fallopian tube, motile cilia are especially important in capturing and initiating oocyte transport in the infundibulum region near the ovary^67–69^. To test the functionality of the cilia induced by β-estradiol treatment and demonstrate the multi-compartment fallopian tube organoid’s mechanical function, we developed a bead displacement assay where oocyte-mimicking fluorescent microbeads were mixed in the lumen-like region of organoids that were stimulated with β-estradiol (Fig. 3D). After 7 days of growth, when the organoids had matured, microbeads were tracked using time-lapse confocal microscopy (Fig. 3E, Movie S1). The trajectories of beads were significantly more irregular and covered larger distances when organoids were treated with β-estradiol (Fig. 3F). The stimulated trajectories were much larger than cell movement within the organoids or the system noise without any cells, and followed the boundary of the folded organoid epithelium. This was quantitatively verified by calculating the mean squared displacement (MSD)^70^ of the microbeads over 8 h (Fig. 3G). In the β-estradiol treated condition, microbead velocity was also increased (Fig. 3H), and while there was no significant difference in persistence of microbead movement (persistence time), the total microbead diffusivity was increased in the β-estradiol treated condition as well (Fig. S3C).

The RT-qPCR results had previously indicated an increase in ciliated cells when these organoids were treated with β-estradiol (Fig. 3B). Even so, organoids were digested to confirm that the increase in microbead movement could be due to motile cilia. β-estradiol treated and untreated organoids were digested, cells were isolated from organoid debris, and cell motility in DPBS was tracked (Fig. 3I, Movie S2). If cells had motile cilia, the cilia acted as a motor and the cells moved in DPBS suspension^71^ (Fig. 3J). The cells isolated from β-estradiol treated organoids had higher MSDs than cells isolated from the untreated organoids (Fig. 3K). This was secondary confirmation that cilia move microbeads in the cores of β-estradiol stimulated organoids, thus the multi-compartment organoids capture the organ’s mechanical function in addition to the molecular function measured via hormone stimulation.

The same bead displacement assay was performed in standard organoids (Fig. S3D). Standard organoids grow from a single parent cell where a lumen forms in the necrotic center of an individual organoid^27^. Due to the nature of organoid maturation in the standard model, most standard organoids did not encase microbeads in the organoid lumen. Unlike the multi-compartment model where microbeads were dispersed along the epithelial-lumen interface (Fig. 3E), most beads were not accessible by lumen-facing epithelial cells in standard organoids (Fig. S3E). However, our electron microscopy results revealed that standard organoids form cilia on the lumen and external epithelial surfaces (Fig. S2D), so microbeads in contact with any surface of the organoid could still be moved by cilia if the cilia were motile. The qualitative trajectories of beads tracked in the standard organoid bead displacement assay revealed larger trajectories in the untreated organoids than in those treated with β-estradiol (Fig. S3F). The quantification of these trajectories also showed little difference in the microbead velocities in any experimental group (Fig. S3G); although, the microbead MSDs in untreated organoids were increased compared to the noise of the system (Fig. S3H). There was no statistically significant difference in the persistence time or total diffusivity for the microbeads tracked in these experimental groups (Fig. S3I). Together, these results show that bead movement in standard organoids was more likely due to movement of cells than motile cilia induced by hormone stimulation.

These functional assessments demonstrate that the multi-compartment organoids resemble the fallopian tube organ in both hormone response and transport ability, thus displaying functional similarity to the organ.

### CODA provides architectural feedback for tuning of future organoid iterations

To validate the architecture of the multi-compartment organoid model, we developed a platform that rigorously compared our multi-compartment organoids to a whole healthy fallopian tube using CODA (Fig. 4A). CODA is a deep learning-based pipeline that 3D reconstructs bulk tissues at cellular resolution from serial histological sections^30^. Here, CODA generated a 3D reference map of a whole fallopian tube received from a transplantation center from a donor who had no history of disease. Using this reference map, architectural parameters including the lumen volume and cell concentrations were quantified and then used to meticulously improve our organoid model to best mimic the tissue architecture. Organoid parameters were adjusted iteratively until this quantitative biomimetic comparison revealed marked improvement in structural biomimetic properties of our organoid model, i.e., most quantified parameters were within 25% deviation from the tissue.

**Fig. 4.**
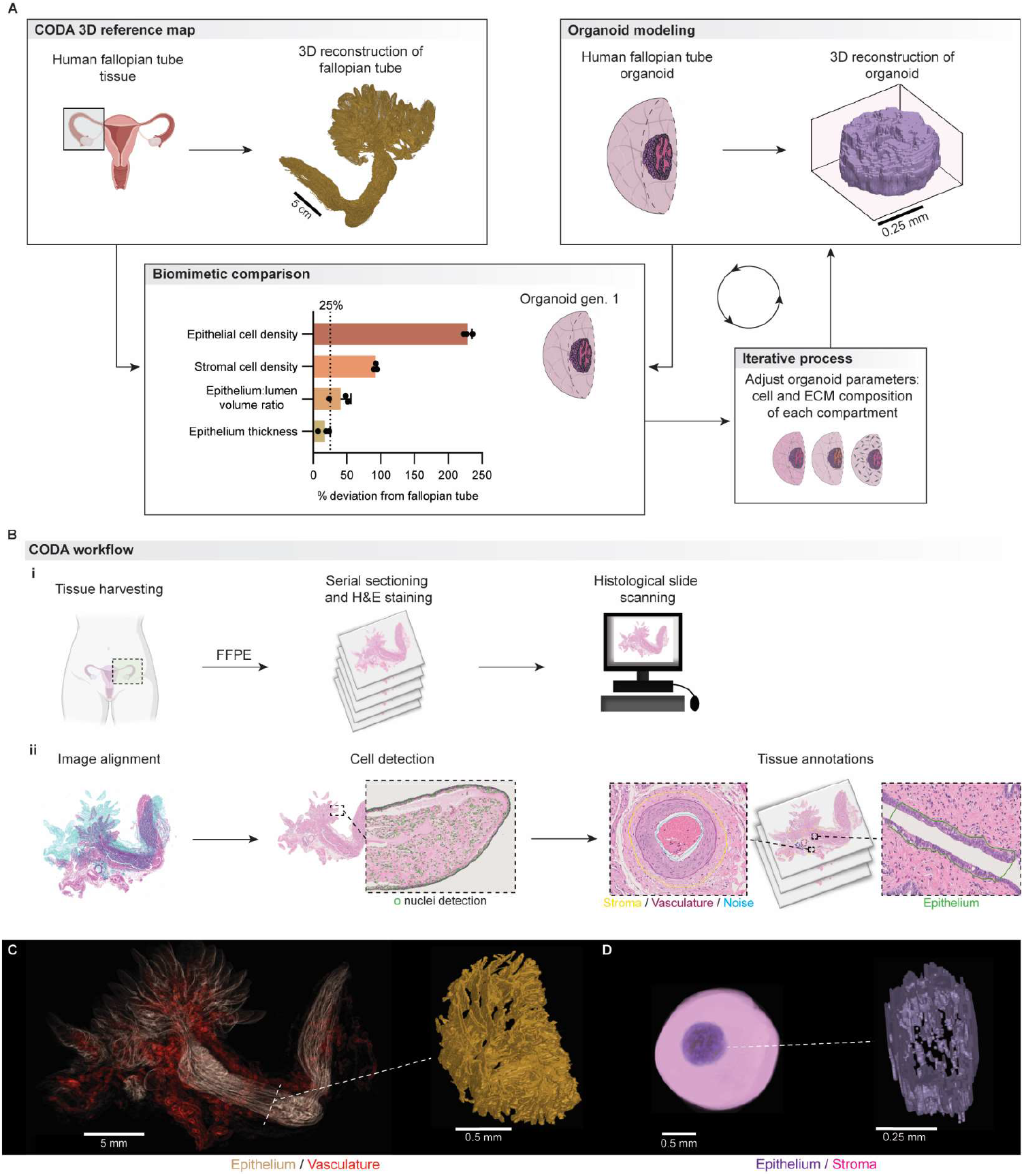
CODA and organoid modeling in tandem produce a tissue-validated organoid model. **A**, An iterative platform compares tissue reference map (top left) and organoid maps (top right). (Bottom left) Architectural parameters were quantified and compared from these maps. (Bottom right) Organoid parameters were adjusted and CODA imaging repeated until an organoid generation closely matches the tissue architecture. **B**, The CODA workflow. (i) A whole fallopian tube was collected and serial sectioned. A subset of the tissue sections was H&E stained. (ii) Sequential tissue sections were aligned to alleviate tissue processing artifacts. Individual cells were detected by their nuclei, and tissue components were annotated in a subset of images. CODA then computationally reconstructs the tissue in 3D. **C**, Best projection of fallopian tube epithelium (gold) and vasculature (red) from a reference map. Scale bar, 5 mm. A 3D reconstruction of a cross section of the fallopian tube is shown (right). Scale bar, 0.5 mm. **D**, Best projection of the organoid epithelium (purple) and stroma (pink) from an organoid map. Multi-compartment organoid parameters: 1 × 10^4^ FTECs in a growth factor reduced Matrigel core, 2 mg/mL collagen I corona. Scale bar, 0.5 mm. A 3D reconstruction of the organoid epithelium is shown (right). Scale bar, 0.25 mm.

The CODA workflow was first applied to reconstruct the fallopian tube of a post-menopausal (72 years old) female donor (Fig. 4B, Fig. S4). Briefly, the sample was formalin fixed and paraffin embedded, serially sectioned (4 µm thickness), and every other section was stained with H&E and digitized for an axial resolution of 8 micron. We registered each H&E image using nonlinear color image registration^30^. Nuclear coordinates were generated through color deconvolution and detection of hematoxylin maxima, and validated through comparison to manually generated coordinates^30^. Fallopian tube epithelium, stroma, vasculature, nerves, and additional structures were labelled at a resolution of 1 micron using a semantic segmentation algorithm. This fallopian tube was assessed by our pathologist investigator to ensure there were no pathological abnormalities and tissue labeling was accurate. Combination of registration, cell detection and tissue segmentation allowed for the production of 3D single-cell resolution maps. We first produced a reference map of a whole healthy human fallopian tube that allowed us to visualize the intricate folding architecture of the epithelium-lumen interface in three dimensions (Fig. 4C).

The same CODA method was applied to our organoids on day 10 of culture for a rigorous side-by-side comparison (Fig. 4A) to 3D reconstruct and label the organoid epithelium and stroma (Fig. 4D). We quantified cellular and extracellular architectural parameters computed from the tissue and the organoid maps to comprehensively compare their structures. We analyzed the level of similarity between organ and organoids, adjusted organoid parameters (ECM composition, cell seeding densities, addition of gynecologic stromal cells), and repeated this process to produce a second iteration (Fig. 4A). Additional organoid parameters were adjusted as necessary to design the next organoid iteration. This iterative process concluded when at least three of four architectural measurements of the organoid were within 25% deviation from the fallopian tube organ (Fig. 4A).

This combined CODA-organoid platform demonstrates a novel feedback platform that combines advanced computational 3D mapping methods and advanced organoid modeling to produce organoids that are micro-anatomically correct. It takes advantage of the customizability and versatility of our multi-compartment organoid model. The result is a precision-engineered organoid that closely resembles the reference fallopian tube. CODA and our organoids are highly versatile and can be applied to any organ/disease, so future organoid models could be analyzed using this platform to examine their architectural and cellular similarity to a healthy or diseased tissue reference map.

### Iterative changes to organoid parameters converge at a next-generation organoid that mimics the tissue architecture

Seven different permutations of organoid epithelial cell seeding densities, organoid core Matrigel composition, and collagen density in the organoid corona were tested in addition to the standard organoid model (Fig. 5A, Fig. S5A). These organoids were not stimulated with β-estradiol to best reflect the post-menopausal conditions of the reference fallopian tube donor, where estrogen levels are low^52,72,73^ and fewer epithelial cells are ciliated^24,52,56^. Hormone stimulation can induce cilia formation (Fig. 3) and could be an additional organoid modification included in future pre-menopausal studies where ciliated epithelial cells are more frequent^24,52,56^ and physiological hormone levels are increased^72^. Beginning with (1) the original organoid parameters (1 × 10^4^ epithelial cells seeded in a growth factor reduced Matrigel core and embedded in a 2 mg/mL collagen I corona), we first increased the collagen density in the organoid corona (2) to 4 mg/mL and 6 mg/mL. Increasing the collagen density reduced the effect of epithelial cell contractility on shrinking the total size of the organoid core, but did not significantly change the epithelial structure within the organoid core (Fig. 5A). We similarly changed the composition of the organoid core ECM by seeding epithelial cells in a Matrigel matrix containing growth factors (3). This adjustment led to a change in the epithelial architecture where epithelial structures remained unconnected within the organoid core, as visualized by confocal microscopy. A similar difference in organoid architecture was also observed when (4) decreasing the epithelial seeding in the organoid core (5 × 10^3^ cells seeded in a growth factor reduced Matrigel core and 2 mg/mL collagen corona).

**Fig. 5.**
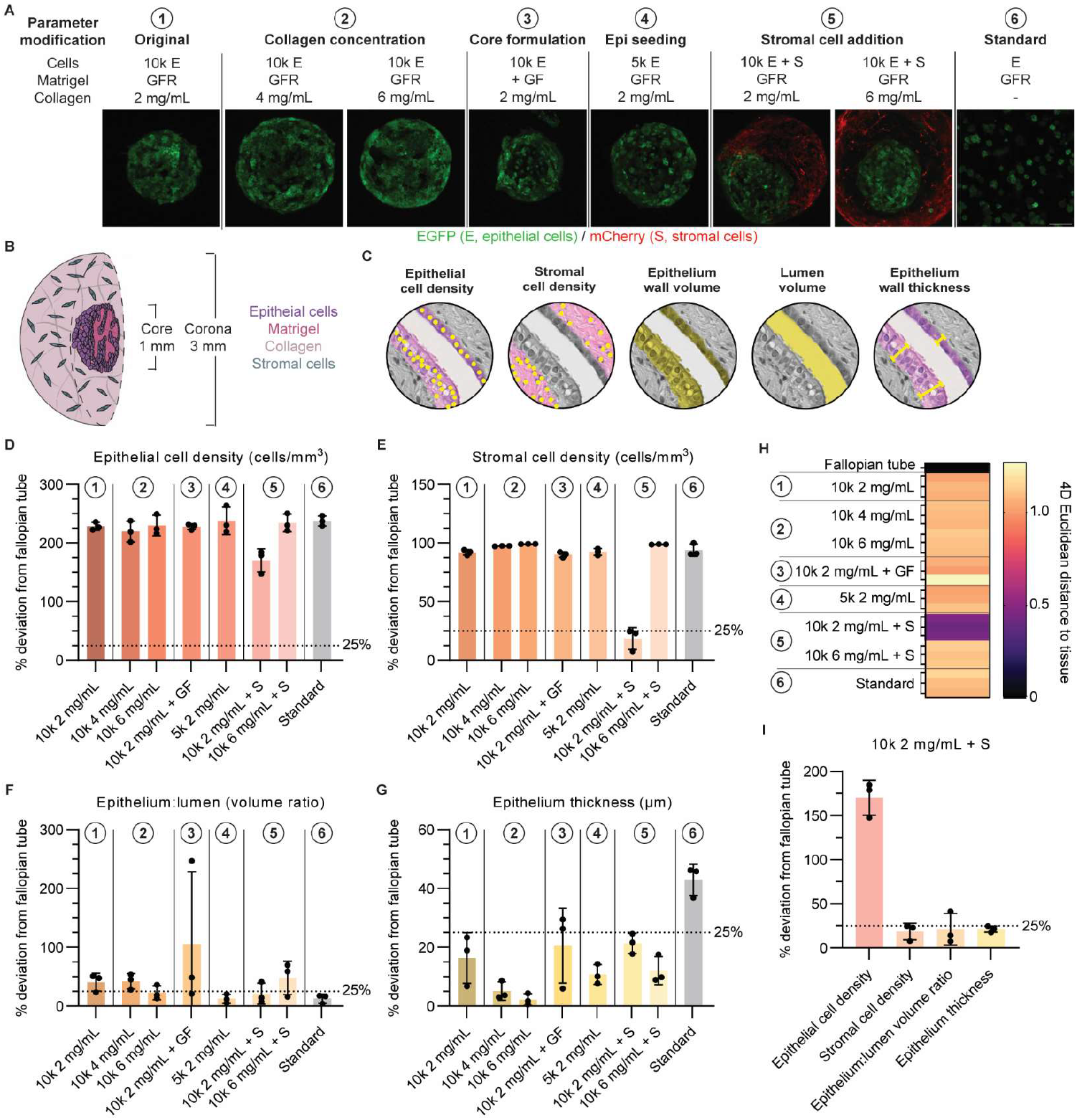
Intentional tuning of organoid parameters produces a model that mimics the fallopian tube. **A**, Cell seeding numbers (10k = 1 × 10^4^, 5k = 5 × 103), cell types (E = FTECs, S = gynecologic stromal cells), Matrigel type (GFR = growth factor reduced, +GF = formulation with growth factors), and collagen density (2, 4, or 6 mg/mL) were adjusted in the multi-compartment model. Lines indicate a new parameter modification. Images are maximum intensity projections. EGFP-tagged FTECs (green) and mCherry-tagged gynecologic stromal cells (red). Scale bar, 250 µm. **B**, Representative cartoon of the multi-compartment organoid with stromal cells in the corona. **C**, Depictions of the architectural parameters quantified from CODA maps. The percent deviation of each organoid generation from the fallopian tube reference values: **D**, epithelial and **E**, stromal cell densities (cells/mm^3^), **F**, volume ratio of epithelium wall to lumen, and **G**, epithelium wall thickness (µm). Lines and numbering indicate a new parameter modification. Dotted line indicates maximum target deviation (25%). n = 3. Data are mean ± SD. **H**, Organoids are quantitatively compared to the healthy fallopian tube via Euclidean distance. **I**, Most architectural measurements are within 25% deviation from the fallopian tube in the final organoid generation.

A unique advantage of the multi-compartment organoid is that stromal cells can be added into the collagen-dense stroma-simulating corona (Fig. 5B). With this modification, we observed significant ECM remodeling when gynecologic stromal cells were seeded in a 2 mg/mL collagen I corona, so we also increased the collagen concentration to 6 mg/mL (Fig. 5A, parameter modification 5). The epithelial architecture, however, was not visibly changed by this addition. The standard organoid technique (6) was also imaged using CODA for a total of eight organoid generations that we analyzed using our new architectural validation platform. Computational analysis was blinded to these experimental conditions.

An array of structural parameters was quantified from each organoid map and compared to the CODA quantifications of the whole fallopian tube (Fig. 5C, Fig. S5B). The measurements from this fallopian tube reference map focused on the ampulla region near the fimbriated end where a defined volume could be measured. This platform can be tuned, however, to focus on different tissue regions and different arrays of architectural parameters specific to the study. The epithelial and stromal cell seeding densities are two parameters of the organoids that are easily tunable, but have different impacts on the organoid architecture (Fig. 5D,E). The inclusion of stromal cells specifically had the most significant impact on organoid similarity to the tissue (Fig. 5E), which highlights the necessity of compartmentalization in an organoid. Stromal cells can only be physiologically incorporated into a fallopian tube organoid model when they are adjacent to the epithelium and in direct contact with a collagen I matrix, different from the epithelial ECM. When incorporating stromal cells, we seeded the stromal cells at a ratio of 6 stromal cells to every 10 epithelial cells (6 × 10^3^ stromal cells in the corona and 1 × 10^4^ epithelial cells in the core) to account for the different proliferation rates of the cell types and the ratios of stromal cells to epithelial cells that had been counted in the tissue reference map (Fig. S5B). The proliferation rate of the stromal cells was further modulated by collagen density. By increasing the collagen density from 2 mg/mL to 6 mg/mL, we limited the proliferation rate of stromal cells, which ultimately decreased the similarity of the stromal cell density per mm^3^ total volume of our organoid on day 10 to the stromal cell density of the tissue within a 1 mm^3^ region surrounding the fallopian tube lumen (Fig. 5E, Fig. S5B). Incorporating stromal cells into the organoid corona also limited the proliferation of the organoid epithelium (Fig. S5C). Additional quantified parameters included the epithelial wall volume and lumen volume, which we reported as a ratio to demonstrate the similarity of the relative scale of the organoid to the relative scale of the reference fallopian tube (Fig. 5F, Fig. S5B). We also measured the thickness of the epithelium wall, which is an indicator of cellular organization (Fig. 5G, Fig. S5B). As shown in Fig. 1A, the fallopian tube epithelium can be a mixture of monolayered regions and regions where the epithelium becomes multi-layered. We used epithelium wall thickness to determine the average epithelium thickness across the whole organ epithelium and confirm that the organoid epithelium organized similarly.

These measured parameters were integrated into Euclidean distances which give equal weights to all metrics (Fig. 5H) and UMAP (Fig. S5D) analyses to compare the fallopian tube tissue with the different organoids. These analyses revealed that the combination of seeding 1 × 10^4^ FTECs in a growth factor reduced Matrigel core and embedding that core in a 2 mg/mL collagen I corona with 6 × 10^3^ stromal cells (10k 2 mg/mL + S) best mimicked the reference fallopian tube tissue. This combination of multi-compartment parameters was a closer reflection of the fallopian tube architecture than the standard organoid. The deviation between this organoid and the reference fallopian tube was < 25% for the stromal cell density, the epithelium-to-lumen volume ratio, and epithelium thickness (Fig. 5I), thus this generation of multi-compartment organoid achieved the target similarity to the reference fallopian tube.

## Discussion

We demonstrate a hybrid CODA-informed organoid engineering platform for the development and validation of organoid models with direct comparisons between the organoid and the organ. This dual protocol can be applied to any organoid model of any tissue/organ and adapted to match the architectural, molecular, and functional parameters of the organ of interest. CODA-based analysis with multi-compartment organoid modeling pair to form a robust platform to develop fine-tuned physiologically and anatomically accurate *in vitro* models for study of the intricacies of many diseases and conditions. Current morphometric-based CODA can be similarly combined with molecular features derived from single cell and spatial transcriptomics to provide improved diagnostics or assessments of disease models.

We present this platform in the context of a healthy human fallopian tube as a basis for future studies of gynecologic disease, such as HGSC^8,13,66,74^. Our multi-compartment organoids of the human fallopian tube surpass the standard organoid model^27,28^ in traditional *in vitro* assessments. Tissue molecular expression patterns are preserved in the multi-compartment organoid, secretory and ciliated epithelial cells are polarized to face the lumen-like region of the organoid, and the response to menstrual cycle hormones (β-estradiol and progesterone) mimics physiological expectations. We show that beyond these established molecular and functional (hormone response^51^) assessments, the mechanical function of the fallopian tube (transport of an oocyte^1,2,67–69^) can also be recreated in the multi-compartment model, but not in the standard fallopian tube organoid via a novel functional assay.

Precision-engineering the organoid model with reference to the architecture of a whole, transplantation-quality human fallopian tube provides an additional quantitative validation of the multi-compartment organoid model. As tunable parameters of the organoid model are modulated, we use CODA imaging and architectural quantification to identify an optimized combination of organoid parameters. While the standard model performed similarly to some multi-compartment configurations, the multi-compartment organoid combination of 6 × 10^3^ stromal cells in a 2 mg/mL collagen I corona with 1 × 10^4^ epithelial cells in growth factor reduced Matrigel (10k 2 mg/mL + S) closely mimicked the fallopian tube tissue’s cellular and extracellular architecture (< 25% deviation for most architectural measurements). The addition of stromal cells in the organoid corona was most significant in achieving architectural similarity of the organoid to fallopian tube tissue. Stromal cells can only be physiologically added to such organoid models when isolated from epithelial cells in a collagen-rich ECM which necessitates the multi-compartment model.

Our multi-compartment organoid model of the human fallopian tube is a tissue-validated healthy model that can be tuned to reflect the diseased states of the fallopian tube in future studies investigating early HGSC or other gynecologic conditions involving the fallopian tube. The cell-cell and cell-ECM interactions in the organoid model can be customized to reflect diseased tissues and referenced to corresponding CODA maps via this iterative platform. We focus on the mechanical cell-ECM interactions and paracrine signaling between epithelial and stromal populations separated into two organoid compartments to maintain a physiological microenvironment. The versatility of this approach provides the opportunity to include additional cell mechanical interactions or introduce disease-related ECM changes in downstream experimental applications.

## Acknowledgements

The authors would like to thank all members of the Wirtz Lab for their feedback. We also thank Mark Atkinson, Mingder Yang, Robert O’Flynn and the team at the Network for Pancreatic Organ Donors with Diabetes (nPOD) for coordinating tissue samples. We thank Ronny Drapkin and his lab for the gifted FTEC cell lines. This work also would not have been possible without the help of the Sidney Kimmel Comprehensive Cancer Center Oncology Tissue Services at Johns Hopkins and the Johns Hopkins School of Medicine Microscope Core Facility. This work was supported through grants from the National Cancer Institute (U54CA143868 and U54CA268083 to D.W.), the National Institute of Arthritis and Musculoskeletal and Skin Diseases (U54AR081774 to D.W.), the National Institute on Aging (U01AG060903 to D.W.), and the Ovarian Cancer Specialized Program of Research Excellence (SPORE) at Johns Hopkins University.

## Author contributions

A.J.C., A.F., and D.W. developed the hypothesis and designed the experiments. A.J.C. performed wet lab experiments and data analysis assisted by I.B., T.R., D.S., K.R., J.B., W.H., G.C.R., and M.H.L. The dry lab experiments and data analysis were performed by A.F. assisted by V.Q., D.K., and A.K. Histological tissue analysis was verified by I.M.S. and T.L.W. Image analysis was performed by P.H.W. The manuscript was written by A.J.C., D.W., A.F., and A.K. with input from I.M.S., T.L.W., and B.S.

## Data and materials availability

All data are presented in the main or supplementary materials. All raw data, cell lines, and MATLAB codes are available upon request.

## Supplementary Materials

Materials and Methods

Figs. S1 to S5

Tables S1 to S2

Supplementary References

Movies S1 to S2

