## Supplementary Materials for "Precision-engineered biomimetics: the human fallopian tube"

### Materials and Methods

#### *Fallopian tube tissue*

Whole healthy fallopian tube tissues were received from organ procurement organizations from donors with no history of disease. All procedures were approved by the Johns Hopkins Internal Review Board (IRB00324065). The tissues were transported in saline on wet ice, formalin fixed (10%, VWR 16004-121) immediately and paraffin embedded. The entire tissue was serial sectioned at 4 µm thickness at Johns Hopkins Oncology Tissue Services (SKCC). Tissue sections were stained with hematoxylin and eosin, or immunohistochemistry (IHC) stained. Antibodies used in IHC staining are listed in Table S1. Tissue sections were scanned at 20X or 40X magnification (Hamamatsu NanoZoomer S210).

#### *Cell culture*

Fallopian tube epithelial and stromal immortalized cells were the FT2821J[1–4], FT194 (CVCL_A4AW)[2] and hS1[1] cell lines. Cells were cultured in RPMI 1640 (Gibco 11875-093) with 10% FBS (Corning 35-010-CV) and 1% penicillin-streptomycin (Sigma P0781). Fallopian tube primary epithelial cells (LifeLine Cell Technology FC-0081) were cultured in complete ReproLife$^{TM}$ Reproductive Medium (LifeLine Cell Technology LL-0068) for up to 3 passages. All cells were cultured at 37°C and 5% $CO_2$ in a humidified incubator and did not surpass 20 cell passages.

Fluorescently tagged cells were transduced via lentivirus delivery of the fluorescent tag. Luciferase expressing cells were transduced via lentivirus delivery of a luciferase-mCherry tag (pSLCAR-CD19-BBz [Addgene 135992] with luciferase [Addgene 46793] and mCherry [Addgene 84020] sequences inserted). Transduced populations were expanded and sorted at ≥ 95% positivity by fluorescence-activated cell sorting (SONY SH800).

#### *Multi-compartment organoid*

The protocol for generating multi-compartment organoids is described in greater detail by Lee, *et al.,* where the protocol was first established as a tumor organoid model[5]. In short, a single-cell suspension of 1 x $10^4$ or 5 x $10^3$ fallopian tube epithelial cells per µL Matrigel (standard formulation 354234 or growth factor reduced 354230, Corning) were pipetted in 1 µL volumes in columns (USA Scientific 1111-3700) containing mineral oil (Sigma 69794) and allowed to gel at 37°C. After 1 h, these cores were collected in cell culture medium (RPMI 1640, Gibco 11875-093) to wash off the mineral oil and embedded in collagen I (high concentration rat tail type I, Corning 354249). A final concentration of 2, 4 or 6 mg/mL collagen I was used according to the protocol previously described[6–9]. Drops containing one organoid core and 10 µL of collagen I were pipetted into fresh mineral oil columns and again allowed to gel at 37°C. After 1.5 h, these organoids were collected in cell culture medium to wash off any remaining mineral oil and plated in a 96-well round bottom plate (Genesee Scientific 25-224) with normal cell culture medium as described above. All organoids were cultured at 37°C and 5% $CO_2$ in a humidified incubator. These multi-compartment organoids are medium throughput, generating up to 96 organoids per batch.

#### *Standard organoid*

The protocol for generating standard organoid is described in greater detail by Kessler, *et al.*[10] Briefly, fallopian tube epithelial cells were suspended in growth factor reduced Matrigel (Corning 354230) at 500 cells/µL. 50µL drops of the mixture were plated on a glass bottom plate and allowed to gel. Organoids were grown in organoid formation medium containing Advanced DMEM/F-12 (Gibco 12634010), L-Glutamine (1%, Gibco 25030081), B-27 (2%, Gibco 17504044), EGF (0.1 µg/mL, ThermoFisher PHG0311), Y-27632 ROCK Inhibitor (10 µM, Cayman Chemical 10005583), TGF-β RI Kinase Inhibitor VI SB431542 (0.5 µM, Sigma 616464) and HEPES (1%, Gibco 15630080). All organoids were cultured at 37°C and 5% $CO_2$ in a humidified incubator.

#### *PrestoBlue proliferation assay*

Organoid total proliferation was assessed via a PrestoBlue (ThermoFisher A13262) proliferation assay. Organoids were incubated with PrestoBlue (10X) diluted to 1X in culture medium for 3 hours at 37°C and 5% $CO_2$ in a humidified incubator. Red fluorescence (RFU) was read on a SpectraMax plate reader (Molecular

Devices, excitation 540, emission 600, cutoff 590). The average of background wells of 1X PrestoBlue in medium without organoids was subtracted from all RFU values. PrestoBlue solution was washed from organoids by replenishing culture medium twice with DPBS (Corning 21-031-CV) and twice with culture medium, and cultured until the next timepoint at 37°C and 5% $CO_2$ in a humidified incubator.

#### *Organoid imaging*

Organoids were imaged on a Nikon Ti2 microscope (phase-contrast) and a Nikon A1R confocal/Nikon Eclipse Ti inverted microscope (DIC, fluorescence and immunofluorescence confocal). The Nikon Ti2 microscope was equipped with a DS-Qi2 camera and a 10X/0.30 Plan Fluor Ph1 DL objective (Nikon). Temperature and $CO_2$ were controlled with a Tokai Hit stage top incubator. The Nikon A1R confocal/Nikon Eclipse Ti inverted microscope was equipped with a 10X/0.30 DIC L/N1 and 20X/10.75 DIC M/N2 Plan Fluor objective (Nikon), a 10X and 20X DIC slider (Nikon), and a N1 and N2 DIC condenser (Nikon). Temperature and $CO_2$ were controlled with an OKO Labs stage top incubator.

Fluorescence and immunofluorescence confocal images are maximum intensity projections of z-stacks taken over 200-500 µm total height of the organoid with a 10 µm interval. Organoids were fixed overnight in 4% PFA or grown in a glass bottom plate for imaging (EGFP, mCherry). Nuclei were stained with Hoechst 33342 (ThermoFisher H3570) and F-actin was stained with Phalloidin 488 (Invitrogen A12379) for 1 hour. The primary and secondary antibodies used in immunofluorescence (IF) imaging are outlined in Table S1. For IF studies, organoids were permeabilized in 0.1% Triton X-100 (Sigma T9284) in DPBS (Corning 21-031-CV) for 1 hour, blocked with 5% normal goat serum (NGS, Cell Signaling Technology) in DPBS for 3 hours, and primary stained overnight in 1% NGS in DPBS. Organoids were secondary stained for 3 hours in 1% NGS in DPBS. Organoids were cleared for 20 minutes in a fructose-glycerol clearing solution[11] (60% glycerol [JT Baker 2136-03], 2.5M D-fructose [Sigma F0127] in DI water) and imaged in DPBS or clearing solution.

#### *Transmission electron microscopy*

Organoids were fixed (2.5% glutaraldehyde, 0.1M sodium cacolydate buffer, 3 mM magnesium chloride in DI water), and imaged via transmission electron microscopy at the Johns Hopkins School of Medicine Microscope Facility.

#### *Western blot*

Protein was extracted from organoids and 2D cells with 2X clear sample buffer (120 mM Tris pH 6.8, 4% SDS, 20% glycerol) and 1X protease/phosphatase inhibitor (Cell Signaling Technology 5872). 2D cells were scraped from a 10 cm dish after coating the plate with 2X clear sample buffer. All protein samples were sonicated with a needle probe sonicator (10-15 pulses, VWR Scientific) and denatured at 100°C for 5 minutes. Samples were stored at -80°C. Protein concentration was measured by microBCA (ThermoFisher, 23235) and absorbance measured on a SpectraMax plate reader (Molecular Devices, 562nm). Samples were diluted to 1 µg/µL in 2X Lamelli (BioRad 1610737) and heated for 5 minutes at 100°C. Samples were loaded in a 4-15% mini-protean gel (BioRad 4561086) for gel electrophoresis (BioRad PowerPac). Protein bands were transferred to a PVDF membrane using a Transblot Turbo pack (BioRad 1704156). The blot was blocked in blotting-grade blocker (5%, 1706404) in TBST (1X Tris Buffered Saline [BioRad 1706435], 0.1% Tween 20 [Sigma P1379], in DI water) for 1 hour and primary stained overnight at 4°C. HRP-conjugated secondary antibodies were incubated at room temperature for 1 hour. Western blots were imaged on a BioRad ChemiDoc XRS+ imager with Prometheus™ ProSignal™ HRP substrate (Genesee Scientific 20-302). Antibodies are listed in Table S1.

#### *Hormone stimulation and RT-qPCR*

After 7 days of organoid culture, organoids were treated with β-estradiol (500 pmol/L, Sigma E2758) or progesterone (50 ng/mL, Fisher Scientific AC225650250)[10]. Hormones were diluted in ethanol prior to dilution in culture medium. This dilution was equivalently reflected in the control. Organoids were grown for another 7 days with hormone stimulation before mRNA harvested (Zymo Research R2052 Direct-zol RNA MiniPrep 11-331) and cDNA synthesized using the iScript™ cDNA Synthesis Kit (BioRad 1708891) and a Blue-Ray Biotech thermocycler. Expression of select genes (primers from Integrated DNA Technologies listed in Table S2)[10,12–15] was quantified by iTaq-SYBR Green (BioRad) RT-qPCR on a BioRad CFX384 Touch Real-Time PCR detection system. Relative expression was calculated according to the equations described by Pfaffl[16].

***Bead displacement assay***

In the bead displacement assay, red fluorescent microbeads (Cospheric FMR-1.3 1-5um) were suspended in Matrigel at the start of organoid generation. Both standard and multi-compartment organoids were generated that contained microbeads only and EGFP-tagged cells with microbeads in the Matrigel. The previously outlined protocols for organoid generation were followed; however, the multi-compartment organoids were plated in 100 µL collagen I in a flat glass bottom plate so that the same location on an organoid could be imaged via time-lapse microscopy. Organoids were treated with β-estradiol[17] (10 nM, Sigma E2758) or control medium after 24 hours. After 7 days of culture, organoids were imaged via confocal fluorescence microscopy (10X magnification). Time-lapse images were collected every 10 minutes for 12 hours. Stage drift was removed from time-lapse images and microbeads tracked across all time points. Microbead trajectories, mean squared displacements and motility information was calculated according to previously published protocols[18].

***Cilia-driven motility assay***

Multi-compartment organoids with 10 µL coronas were cultured according to the same specifications of the bead displacement assay (β-estradiol [10 nM, Sigma E2758][17] treated after 24 hours, grown for 7 days) and digested with 2 mg/mL collagenase I (Gibco 17018029) for 1 hour. After digestion, cells were resuspended in DPBS (Corning 21-031-CV) and time-lapse imaged for 1 hour (phase-contrast, 10X magnification). After an initial 30 minute incubation to give cells time to settle to the bottom of the plate, cells were imaged for 30 minutes at 1 minute intervals. Individual cells were tracked and their motility parameters calculated according to previously published cell tracking protocols[18].

***CODA***

The CODA protocol is described in greater detail by Kiemen, *et al.,* where the protocol was first established[19]. Organoids after 10 days of growth and fallopian tube tissues within 12 hours of collection were formalin fixed (10%, VWR 16004-121) and paraffin embedded, then serial sectioned across the whole tissue at 4 µm section thickness. One in every two tissue slides was designated for H&E staining, while the remaining unstained slides were stored for potential application of multiplexing techniques, such as immunofluorescence, immunohistochemical, and tissue-omics. Stained slides were scanned at 20X magnification offering a 0.5 micron resolution on the x and y dimensions. To align the H&E images, a combination of global and nonlinear techniques was implemented in all images in a sample. To statistically measure the similarity between two serially aligned H&E images, cross-correlation was used. In order to identify individual cells on the H&E-stained samples, a color deconvolution step was added which extracted the hematoxylin channel of each image, and specific sized and spatially distanced two dimensional minima were marked as nuclei. A deep learning model was trained on an image subset using manually annotated tissue cases. To enhance the performance of the convolutional neural network on unseen data, data augmentation techniques were integrated, allowing for hue, rotation, and scale changes to the training dataset. The validation of the model was ensured using independent testing images, not used for training, with the mean testing accuracies surpassing ninety percent for true positive segmentation values. Combination of the alignment, semantic segmentation and cell detection computational methods resulted in three dimensional maps of each individual tissue label at single cell resolution. Three-dimensional renderings were performed at 4 µm and 1 µm in *x* and *y* dimensions for the whole human fallopian tube and each individual organoid, respectively. Lastly, three-dimensional analysis of the human fallopian tube and organoids were performed in isotropic dimensions 4x4x4 $\mu m^3$.

***Illustrations***

Illustrations and cartoons were generated using Adobe Illustrator or BioRender.com.

***Statistical analysis***

Statistical analyses are described in the figure legends with P values.

### Supplementary Figures

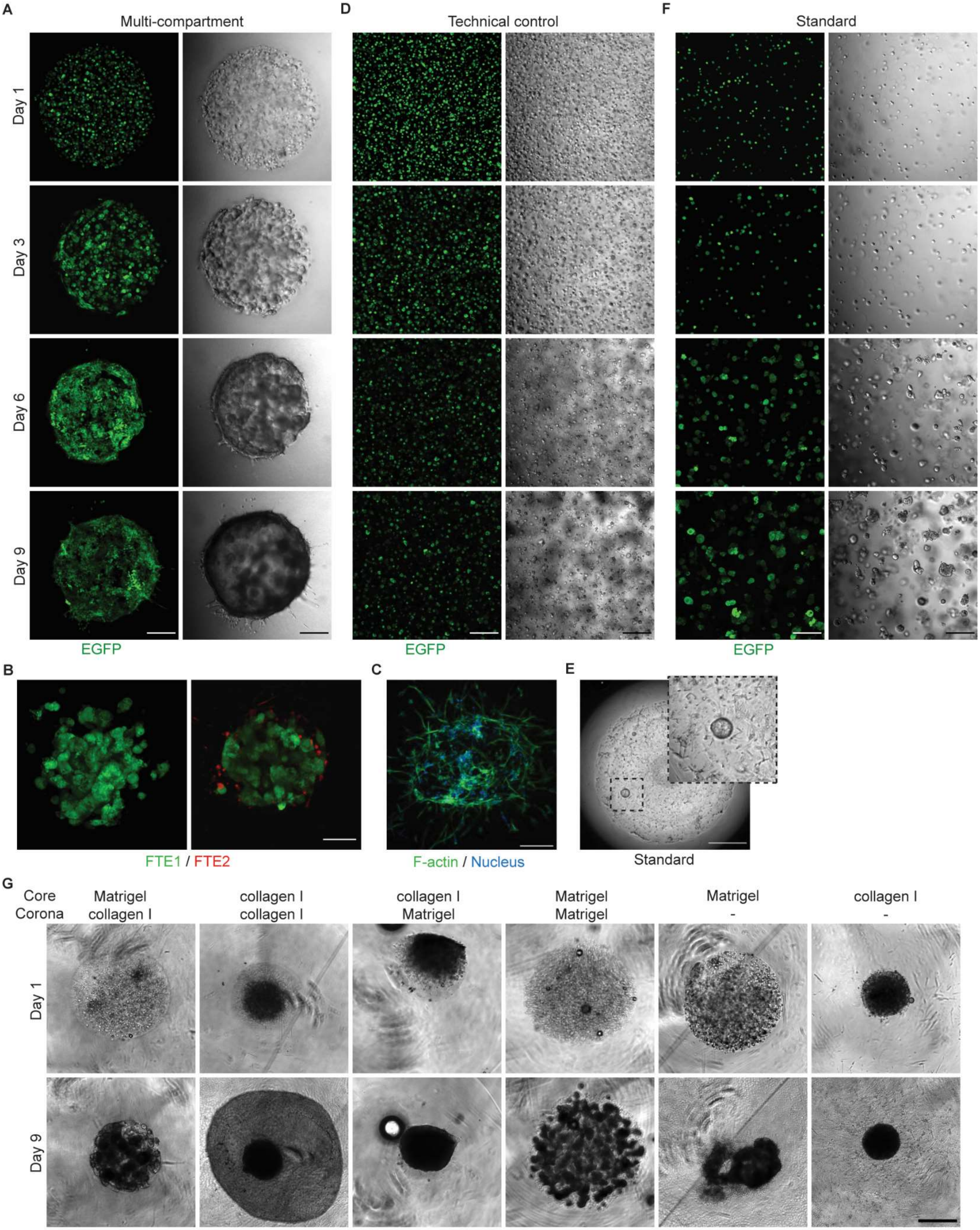

**Fig. S1. Fallopian tube organoid assembly. A**, EGFP-tagged (green) multi-compartment organoid time series. Images are maximum intensity projections (left) and DIC (right). Scale bar, 250 μm. **B**, Two types of FTECs are mixed in the multi-compartment organoid core. FTE cell lines are EGFP (green) and mCherry (red) tagged. Images are maximum intensity projections. Scale bar, 250 μm. **C**, Primary FTECs are grown in the multi-compartment organoid model. Images are maximum intensity projections. F-actin (green) and nuclear DNA (blue). Scale bar, 250 μm. **D**, EGFP-tagged cells (green) are grown at the multi-compartment cell density (1 $x10^4$ cells/μL) and normal cell culture medium, but plated according to the standard organoid technique. Images are maximum intensity projections (left) and DIC (right). Scale bar, 250 μm. **E**, Primary FTECs are grown in the standard organoid model. Image is phase-contrast microscopy. Scale bar, 250 μm. Inset shows one successfully formed standard organoid. **F**, EGFP-tagged (green) standard organoid time series. Images are maximum intensity projections (left) and DIC (right). Scale bar, 250 μm. **G**, Organoids with Matrigel or collagen I in the core and all combinations of corona ECM. Phase-contrast images. Scale bar, 500 μm.

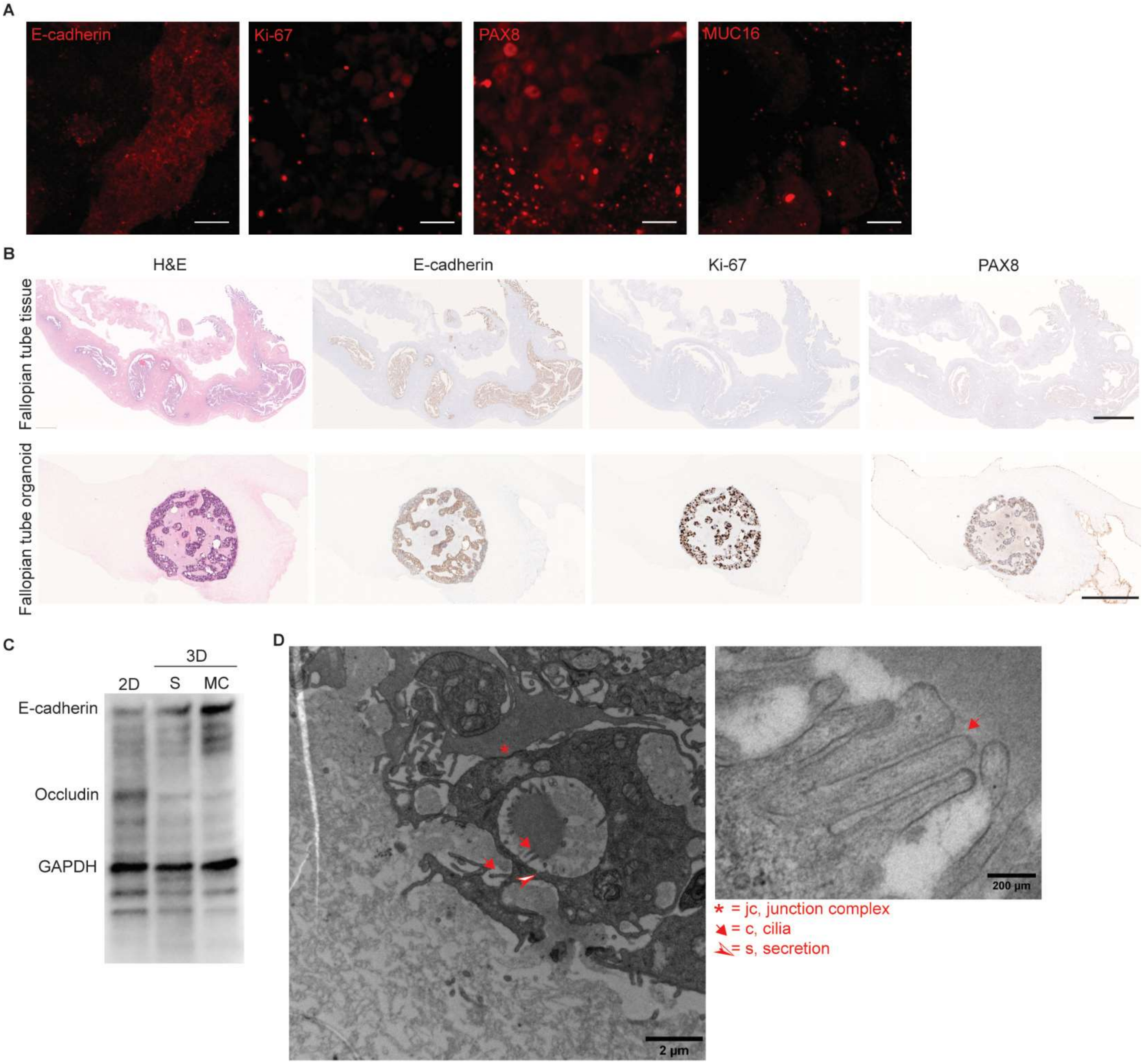


**Fig. S2. Multi-compartment and standard organoid molecular expression patterns and cellular structure. A**, Isolated protein of interest channel for immunofluorescence in multi-compartment fallopian tube organoids. Images of organoids are maximum intensity projections. Scale bar, 20 μm. From left to right: E-cadherin, Ki-67, PAX8, MUC16. **B**, FFPE tissue sections stained with H&E or IHC stained for E-cadherin, Ki-67 and PAX8 (from left to right). Whole, healthy transplantation-quality tissue sections of the fallopian tube (top) and the multi-compartment organoid (bottom). Scale bar, 4 mm (top) and 300 μm (bottom). **C**, Global protein expression in 2D and 3D (S = standard, MC = multi-compartment) culture via western blot. Loading control = GAPDH. **D**, Transmission electron microscopy of a standard fallopian tube organoid. Scale bar, 2 μm. Zoom, 200 nm. Asterisk = junction complex, arrow = cilia, arrowhead = secretion.

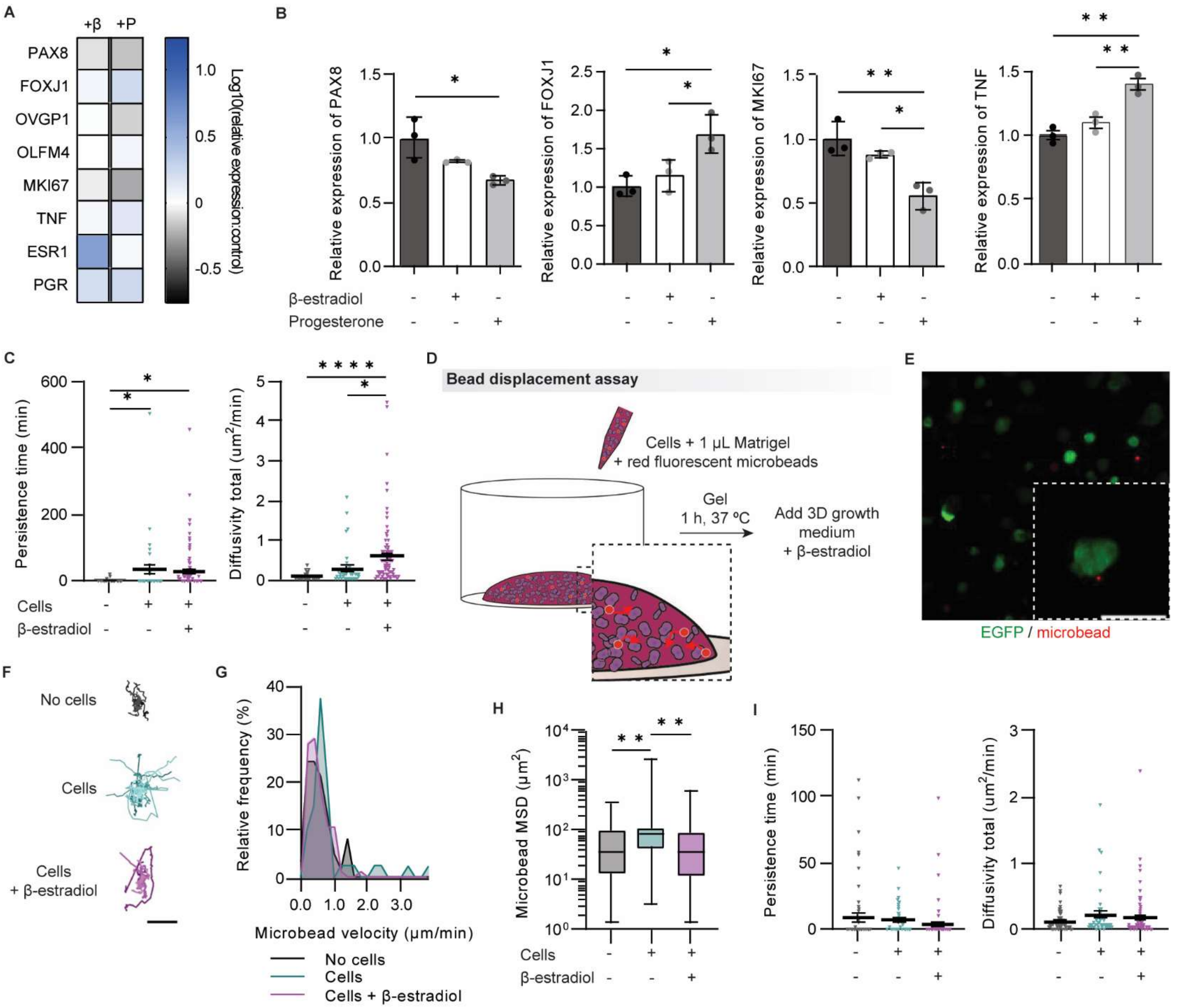


**Fig. S3. Functional response to hormone treatment in organoids.** **A**, RT-qPCR of β-estradiol (β) or progesterone (P) treated standard organoids. Data are log10(relative expression:control). **B**, Relative expression of genes related to cell differentiation into ciliated epithelial cells and proliferation. Data are mean ± SD. **A**-**B**, N = 1, n = 3. **C**, Persistence time and total diffusivity of microbeads in the multi-compartment bead displacement assay. N = 3, n = 4-46. **D**, Cartoon depicting bead displacement assay generation in standard organoids. **E**, A standard organoid used in the bead displacement assay with a microbead at the organoid's epithelial interface. EGFP-tagged FTECs (green) and microbeads (red). Scale bar, 100 μm. **F**, Representative microbead trajectories from standard organoids with microbeads only (noise, black), microbeads with FTECs (teal), and microbeads with FTECs treated with β-estradiol (purple). Scale bar, 7.5 μm. **G**, Relative frequency of microbead velocities in standard organoids. **H**, Microbead MSD over 8 h. **I**, Persistence time and total diffusivity of microbeads in the standard organoid bead displacement assay. **G**-**I**, N = 2, n = 18-45. **C** and **H**-**I**, data are mean ± SEM. Statistical test: **B**-**C** and **H**-**I**, one-way ANOVA, ****$P \leq 0.0001$, **$P \leq 0.01$, *$P \leq 0.05$.

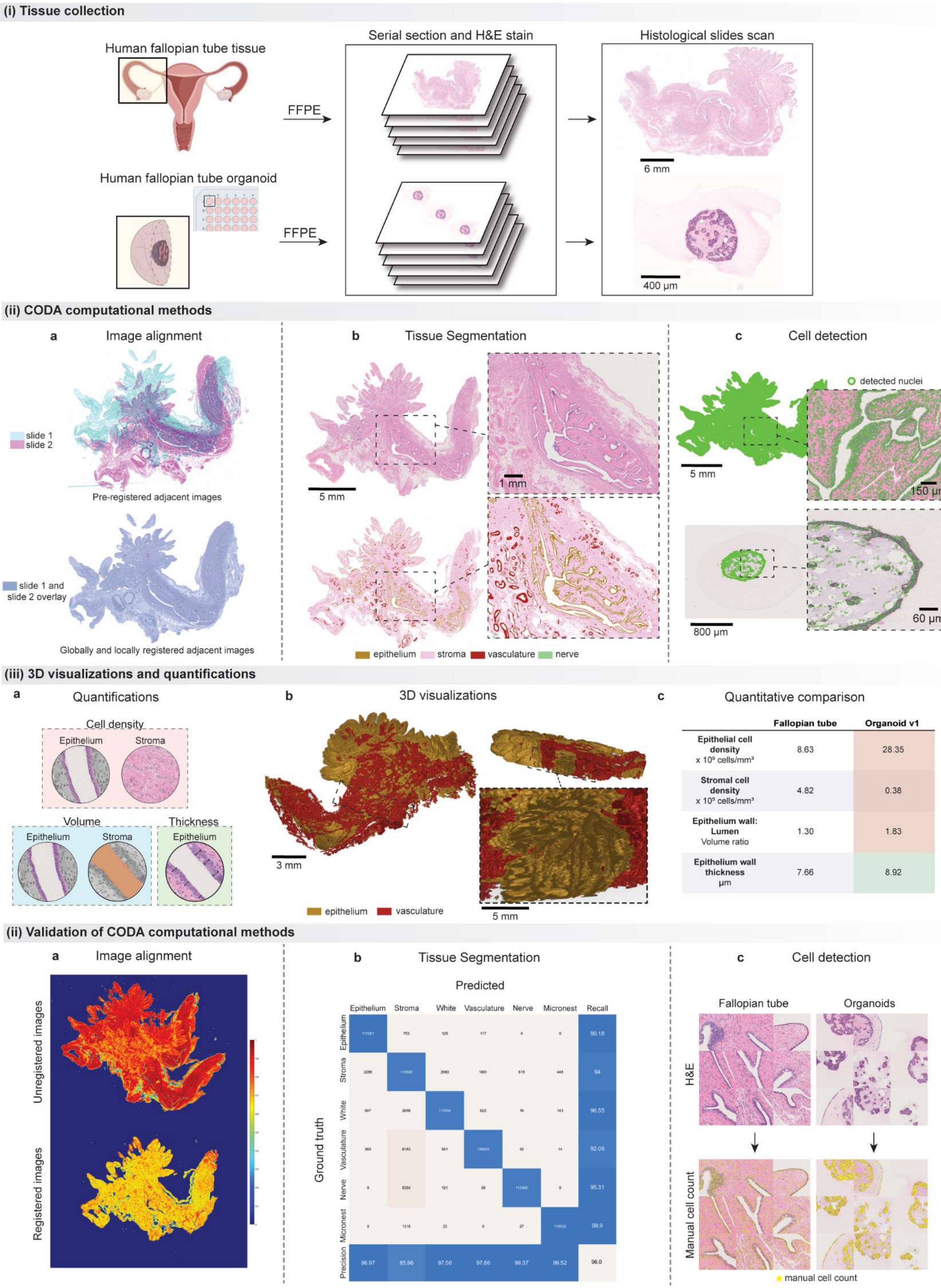

(i) Tissue collection
Human fallopian tube tissue
Human fallopian tube organoid
FFPE
Serial section and H&E stain
Histological slides scan
6 mm
400 µm
(ii) CODA computational methods
a Image alignment
slide 1
slide 2
Pre-registered adjacent images
slide 1 and slide 2 overlay
Globally and locally registered adjacent images
b Tissue Segmentation
5 mm
1 mm
epithelium
stroma
vasculature
nerve
c Cell detection
detected nuclei
5 mm
150 µm
800 µm
60 µm
(iii) 3D visualizations and quantifications
a Quantifications
Cell density
Epithelium
Stroma
Volume
Epithelium
Stroma
Thickness
Epithelium
b 3D visualizations
3 mm
5 mm
epithelium
vasculature
c Quantitative comparison
Fallopian tube
Organoid v1
Epithelial cell density x 10⁵ cells/mm³
8.63
28.35
Stromal cell density x 10⁵ cells/mm³
4.82
0.38
Epithelium wall: Lumen Volume ratio
1.30
1.83
Epithelium wall thickness µm
7.66
8.92
(ii) Validation of CODA computational methods
a Image alignment
Unregistered images
Registered images
b Tissue Segmentation
Predicted
Epithelium
Stroma
White
Vasculature
Nerve
Micronest
Recall
Ground truth
Precision
99.18
94
96.55
92.09
95.31
98.9
96.97
85.96
97.56
97.86
99.37
99.52
96.0
c Cell detection
Fallopian tube
Organoids
H&E
Manual cell count
manual cell count

**Fig. S4. CODA workflow. (i)** Fallopian tube tissue and organoids are formalin fixed and paraffin embedded (FFPE) and the entire tissue is serial sectioned. One in every two tissue sections are stained with hematoxylin and eosin (H&E) and the H&E tissue sections are scanned at 20X magnification. **(ii)** All scanned tissue sections are aligned by an elastic registration process in sequential order. Tissue components such as the epithelium (gold), stroma (pink), vasculature (red) and nerves (green) are manually annotated in a fraction of the images to train the algorithm to segment all tissue sections. Individual cell nuclei are detected from the H&E images. Noise from image scans is adjusted in the cell count. **(iii)** An array of tissue parameters is measured from the processed tissue sections, and fallopian tube tissue and organoids are visualized in 3D. Shown is an epithelium (gold) and vasculature (red) map of the fallopian tube. Architectural parameters quantified from each organoid generation are compared to the tissue. **(iv)** Independent testing of the CODA computational methods validates the performance of our reference maps.

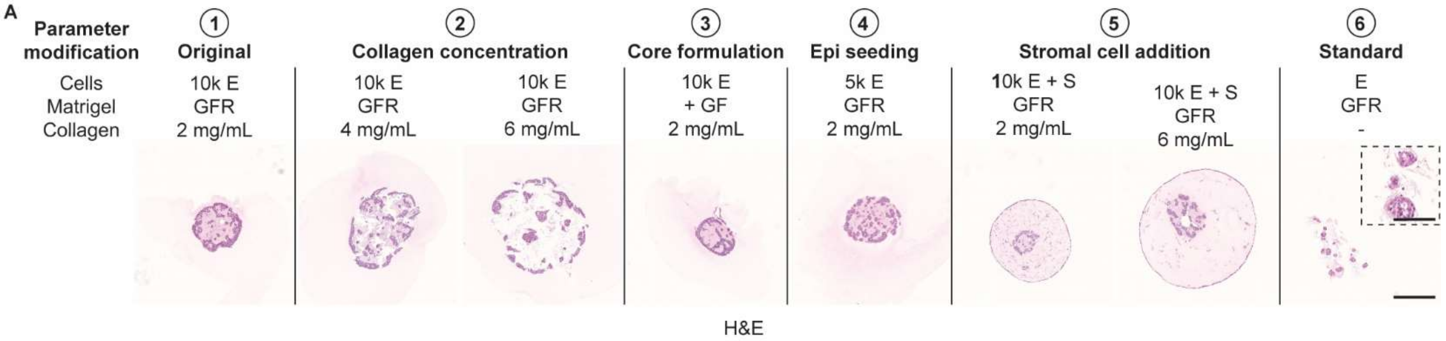


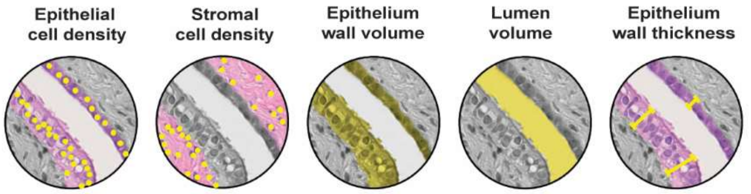


| Iteration | cells/mm³ ( x 10⁵) | cells/mm³ ( x 10⁵) | ratio wall:lumen volume | μm |
|---|---|---|---|---|
| Fallopian tube | 8.63 | 4.82 | 1.30 | 7.66 |
| 10k 2 mg/mL | 28.35 | 0.38 | 1.83 | 8.92 |
| 10k 4 mg/mL | 27.60 | 0.12 | 0.75 | 7.61 |
| 10k 6 mg/mL | 28.47 | 0.04 | 1.26 | 7.72 |
| 10k 2 mg/mL + GF | 28.30 | 0.47 | 2.67 | 9.24 |
| 5k 2 mg/mL | 29.18 | 0.37 | 1.24 | 6.84 |
| 10k 2 mg/mL + S | 23.34 | 3.93 | 1.57 | 6.03 |
| 10k 6 mg/mL + S | 28.85 | 0.06 | 1.26 | 7.63 |
| Standard | 29.14 | 0.29 | 1.13 | 4.37 |

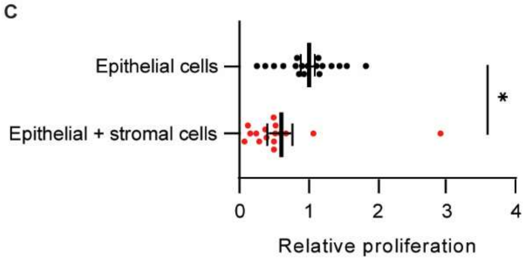


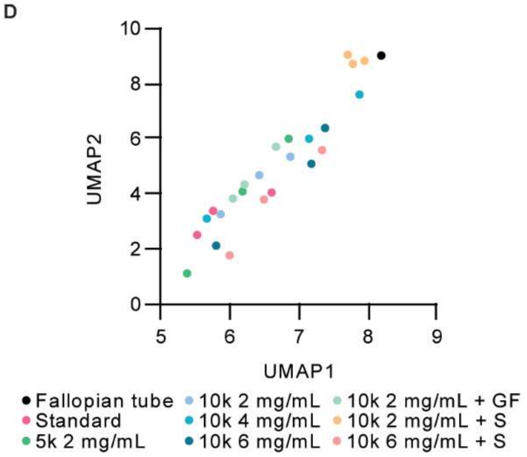


**Fig. S5. Organoid iterations assessed by CODA. A,** H&E stained FFPE tissue sections of all organoid iterations. Cell seeding numbers (10k = $1 \times 10^4$, 5k = $5 \times 10^3$), cell types (E = FTECs, S = gynecologic stromal cells), Matrigel type (GFR = growth factor reduced, +GF = formulation with growth factors), and collagen density (2, 4, or 6 mg/mL) were adjusted in the multi-compartment model. Lines indicate a new parameter modification. H&E: nuclei (purple) and ECM (pink). Scale bar, 500 μm. Inset, 100 μm. **B**, CODA quantified parameters. From left to right, the epithelial and stromal cell densities (x $10^5$ cells/mm$^3$), volume ratio of epithelium wall to lumen, and epithelium wall thickness (μm). Values are the averages of 3 organoids per iteration. **C**, Stromal cells regulate epithelial cell proliferation in the multi-compartment organoid model. FTECs are luciferase-tagged in the organoid core with stromal cells in the corona. Luminescence measured on day 10. N = 3, n = 3-6. Statistical test: unpaired t-test, *$P \leq 0.05$. Data are mean ± SEM. **D**, Organoids are quantitatively compared to the healthy fallopian tube via UMAP analysis. Three organoids per iteration.

### Supplementary Tables

**Table S1.** Antibodies used in immunofluorescence (IF), immunohistochemistry (IHC), and western blots (WB).

| Application | Antibody | Company | Ref / product # |
|---|---|---|---|
| IF | E-cadherin | Cell Signaling Technology | 3195 |
| IF | Ki-67 | Cell Signaling Technology | 9449 |
| IF | PAX8 | Cell Signaling Technology | 59019 |
| IF | MUC16 | Cell Signaling Technology | 29623 |
| IF | Anti-rabbit IgG, Alexa Fluor 568 | Invitrogen | A-11011 |
| IF | Anti-mouse IgG, Alexa Fluor 568 | Invitrogen | A-11004 |
| IHC | E-cadherin | Cell Signaling Technology | 3195 |
| IHC | Ki-67 | Abcam | ab16667 |
| IHC | PAX8 | Cell Signaling Technology | 28556 |
| WB | E-cadherin | Cell Signaling Technology | 3195 |
| WB | Occludin | Cell Signaling Technology | 91131 |
| WB | GAPDH | Cell Signaling Technology | 2118 |
| WB | Anti-rabbit IgG, HRP-linked | Cell Signaling Technology | 7074 |

**Table S2.** Primers used in RT-qPCR.

| Gene | FWD/RVS | Sequence |
|---|---|---|
| PAX8 | FWD | AAGTGCAGCAACCATTCAACC |
| PAX8 | RVS | CTGCTCTGTGAGTCAATGCTTA |
| FOXJ1 | FWD | GGAGGGGACGTAAATCCCTA |
| FOXJ1 | RVS | GGTCCCAGTAGTTCCAGCAA |
| OVGP1 | FWD | TCCCACACATCGTCCAAACAT |
| OVGP1 | RVS | TCCCCATGATGAGCTTCTCTG |
| OLFM4 | FWD | ACTGTCCGAATTGACATCATGG |
| OLFM4 | RVS | TTCTGAGCTTCCACCAAAACTC |
| MKI67 | FWD | AAGCCCTCCAGCTCCTAGTC |
| MKI67 | RVS | TCCGAAGCACCACTTCTTCT |
| TNF | FWD | CCTCTCTCTAATCAGCCCTCTG |
| TNF | RVS | GAGGACCTGGGAGTAGATGAG |
| ESR1 | FWD | CCCACTCAACAGCGTGTCTC |
| ESR1 | RVS | CGTCGATTATCTGAATTTGGCCT |
| PGR | FWD | ACCCGCCCTATCTCAACTACC |
| PGR | RVS | AGGACACCATAATGACAGCCT |
| 18S | FWD | AGAAGTGACGCAGCCCTCTA |
| 18S | RVS | GAGGATGAGGTGGAACGTGT |
| ATUB | FWD | AGGAGTCCAGATCGGCAATG |
| ATUB | RVS | GTCCCCACCACCAATGGTTT |
| ACTB | FWD | CATGTACGTTGCTATCCAGGC |
| ACTB | RVS | CTCCTTAATGTCACGCACGAT |
| GAPDH | FWD | GGAGCGAGATCCCTCCAAAAT |
| GAPDH | RVS | GGCTGTTGTCATACTTCTCATGG |
| RPL13A | FWD | CCTGGAGGAGAAGAGGAAAGAGA |
| RPL13A | RVS | TTGAGGACCTCTGTGTATTTGTCAA |

**Supplementary Movies**

**Movie S1.** An oocyte-mimicking fluorescent microbead is transported along the multi-compartment organoid epithelium in the bead displacement assay. Microbead (red) and EGFP-tagged FTECs (green), treated with β-estradiol. Scale bar, 50 μm.

**Movie S2.** A cell from a β-estradiol-treated organoid moves in DPBS solution after organoid digestion in the cilia-driven motility assay. Scale bar, 50 μm.
